# Structural basis of saccharine derivative inhibition of carbonic anhydrase IX

**DOI:** 10.1101/2023.11.01.565239

**Authors:** Janis Leitans, Andris Kazaks, Janis Bogans, Claudiu T. Supuran, Inara Akopjana, Jekaterina Ivanova, Raivis Zalubovskis, Kaspars Tars

## Abstract

This scientific study explores the binding mechanisms of saccharine derivatives with human carbonic anhydrase IX (hCA IX), an antitumor drug target, with the aim of facilitating the design of potent and selective inhibitors. Through the use of crystallographic analysis, we investigate the structures of hCA IX - saccharine derivative complexes, unveiling their unique binding modes that exhibit both similarities to sulfonamides and distinct orientations of the ligand tail. Our comprehensive structural insights provide information regarding the crucial interactions between the ligands and the protein, shedding light on interactions that dictate inhibitor binding and selectivity. Through a comparative analysis of the binding modes observed in hCA II and hCA IX, isoform-specific interactions are identified, offering promising strategies for the development of isoform-selective inhibitors that specifically target tumor-associated hCA IX. The findings of this study significantly deepen our understanding of the binding mechanisms of hCA inhibitors, laying a solid foundation for the rational design of more effective inhibitors.

## Introduction

Carbonic anhydrases (CAs, EC 4.2.1.1) are a family of enzymes that catalyze the reversible hydration of carbon dioxide. Although CAs play important physiological roles in the body, the activities of certain human CA (hCA) isoforms have been linked to various diseases.^[1,2]^ For example, the widespread hCA II is implicated in glaucoma and acute mountain sickness (AMS) symptoms, while hCA IX is expressed in only a limited number of normal tissues but is highly overexpressed in many solid hypoxic tumors.^[3,4]^ It has been demonstrated that the enzymatic activity of hCA IX promotes tumor growth, whereas its inhibition constitutes a validated strategy to mitigate tumor spread.^[5,6]^ In order to design selective and efficacious inhibitors for hCA IX, it is imperative to acquire a comprehensive understanding of the binding mechanisms between the ligand and the enzyme. X-ray crystallography, as a valuable tool, can greatly facilitate this endeavor.

One class of potent CA inhibitors are sulfonamides, which have been extensively studied. Sulfonamides bind to the enzyme’s active site and block the binding of a water molecule (acting as nucleophile) to the Zn^2+^ ion. The binding mode of sulfonamides has been extensively studied in various CA isoforms and has provided useful insights into the design of novel inhibitors.^[7,8]^ Saccharin, an artificial sweetener, is another well-known CA inhibitor. It has been shown to be an effective inhibitor of hCA IX, with a similar binding mode to sulfonamides. However, due to the difference in their molecular structure, saccharin and sulfonamides result in different binding orientations.^[9]^ This difference can be useful for the design of ligand tail orientation and the development of more effective inhibitors.

Several studies have explored saccharin-sulfonamide derivatives as inhibitors of CA following the initial discovery of saccharin’s inhibitory properties.^[10,11,12,13,14]^ These investigations have demonstrated that compounds incorporating both sulfonamide and saccharin zinc binding groups (ZBGs) tend to exhibit a stronger affinity for binding through the sulfonamide moiety (Figure 1).^[11]^

**Figure 1.**
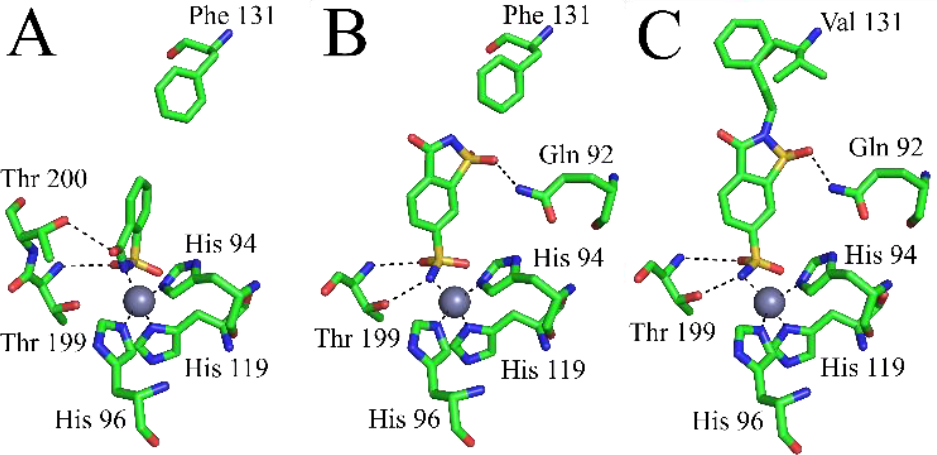
Comparison of binding modes of saccharin (panel A), 6-Sulfamoyl-saccharin (panel B), and compound 1 from this study (panel C). The zinc ion is shown as a gray sphere, His 94, 96 and 119, and residues involved in polar contacts are indicated. Phe/Val 131 is marked to demonstrate the compound’s tail orientation. The figure was prepared using PyMol (DeLano The PyMOL Molecular Graphics System, San Carlos, CA, USA, DeLano Scientific), and the following PDB codes were used: 2Q38 for saccharin, 4XE1 for 6-Sulfamoyl-saccharin and 8Q18 for compound 1.

Compared to structurally similar heterocyclic sulfonamides and acetazolamide (AAZ) (Figure 2), saccharin still exhibits significantly better selectivity towards hCA IX. Inhibition potencies of saccharin and structurally similar heterocyclic sulfonamides and AAZ can be found in Table 1.

**Table 1.** Reported inhibition data of saccharin^[9]^, 6-Sulfamoyl-saccharin^[11]^, hydrochlorothiazide^[9]^, compound 1 from this study^[10]^, AAZ^[9]^ (K_I_ in nM).

|  | hCA I | hCA II | hCA IX |
| --- | --- | --- | --- |
| Saccharin | 18540 | 5950 | 103 |
| 6-Sulfamoyl-saccharin | 3406 | 77 | 12 |
| Hydrochlorothiazide | 328 | 290 | 367 |
| Compound 1 | 9.1 | 0.4 | 378 |
| AAZ | 250 | 12 | 25 |

**Figure 2.**
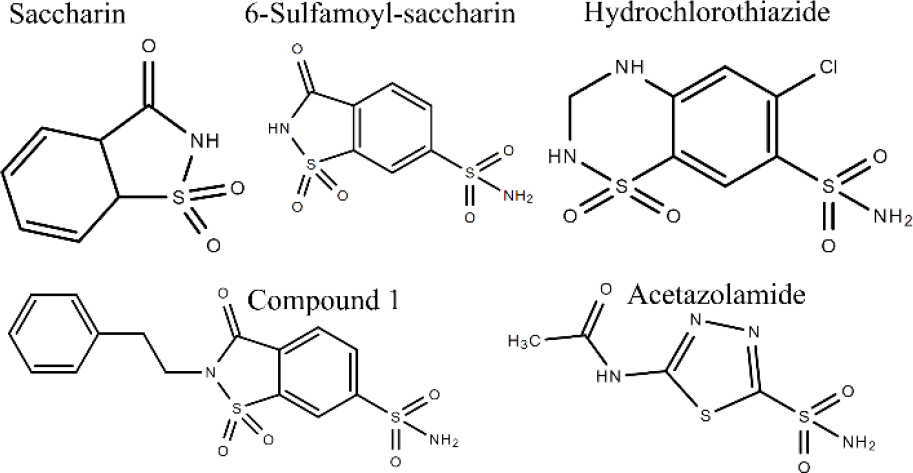
Chemical structures of saccharin, 6-Sulfamoyl-saccharin, hydrochlorothiazide, compound **1** from this study and AAZ.

Our group has previously solved high-resolution crystal structures of human carbonic anhydrase II (hCA II) in complex with saccharine derivatives, revealing interesting binding mechanisms.^[10]^ In this work, we extend this investigation by determining the 3D structures of those compounds, this time in complex with hCA IX. The results of this study will provide valuable insights into the binding mechanisms of these inhibitors with hCA IX and help guide the design of novel inhibitors with increased selectivity and potency.

## Results and Discussion

To gain insights into the binding mechanisms of saccharin sulfonamide derivatives within the active site of human carbonic anhydrase IX (hCA IX), we utilized X-ray crystallography to examine the structures formed between hCA IX and these compounds. While previous studies have investigated the binding mechanisms of these derivatives on hCA II, as described in a published work^[10]^, our focus was specifically on the structures of hCA IX complexed with these compounds. By determining the crystal structures, we obtained detailed information about their precise binding orientations. The resolved structures exhibited resolutions of 2.13 Å for compound 1, 2.63 Å for compound 2, and 2.35 Å for compound 3. A summary of the data processing, refinement, and validation statistics for these structures can be found in Table 2.

**Table 2.**
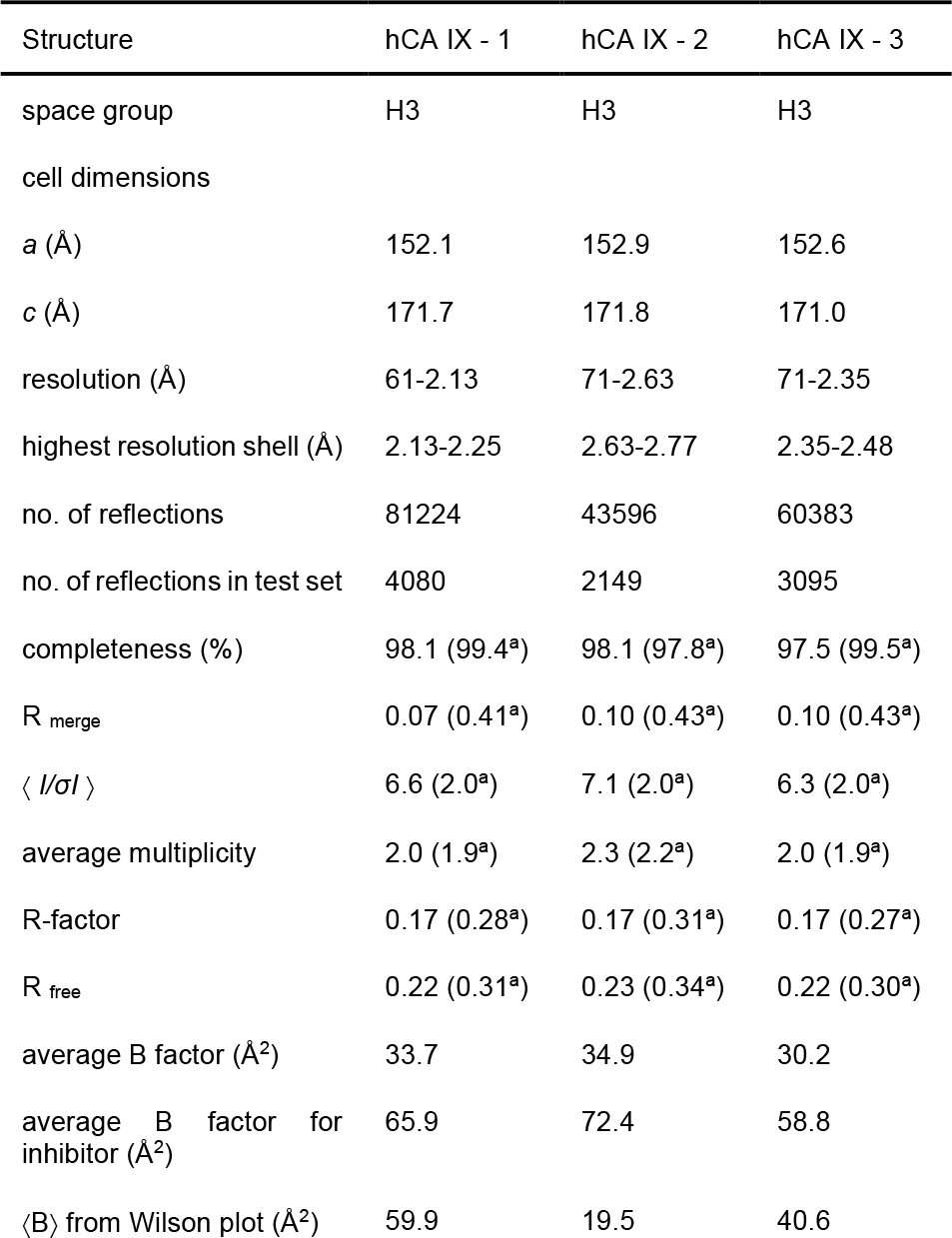
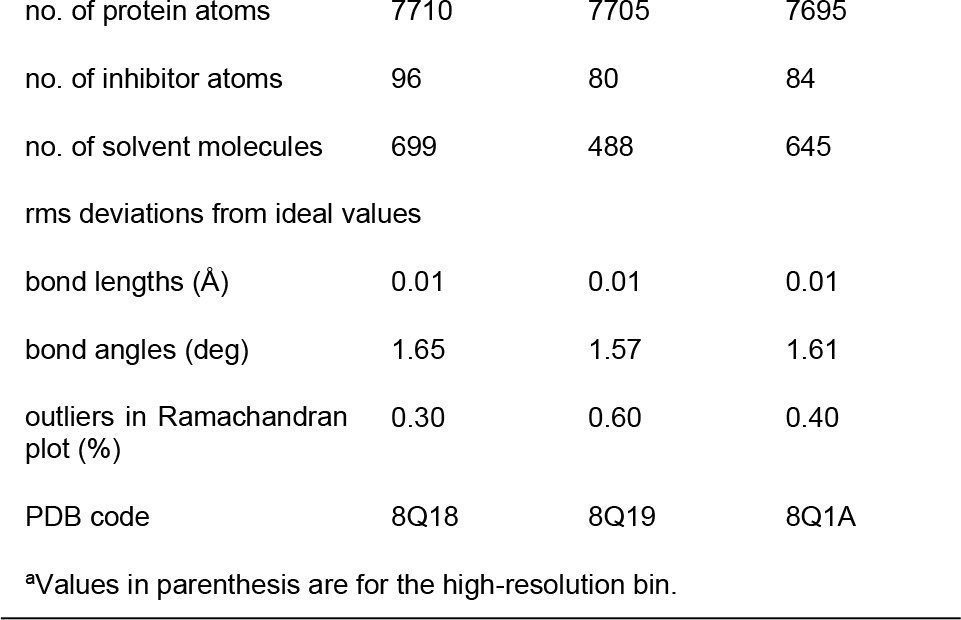
Data processing, refinement and validation statistics for the hCA IX – 1, hCA IX – 2 and hCA IX - 3 Complexes.

| Structure | hCA IX - 1 | hCA IX - 2 | hCA IX - 3 |
| --- | --- | --- | --- |
| space group | H3 | H3 | H3 |
| cell dimensions |  |  |  |
| *a* (Å) | 152.1 | 152.9 | 152.6 |
| *c* (Å) | 171.7 | 171.8 | 171.0 |
| resolution (Å) | 61-2.13 | 71-2.63 | 71-2.35 |
| highest resolution shell (Å) | 2.13-2.25 | 2.63-2.77 | 2.35-2.48 |
| no. of reflections | 81224 | 43596 | 60383 |
| no. of reflections in test set | 4080 | 2149 | 3095 |
| completeness (%) | 98.1 (99.4<sup>a</sup>) | 98.1 (97.8<sup>a</sup>) | 97.5 (99.5<sup>a</sup>) |
| R<sub>merge</sub> | 0.07 (0.41<sup>a</sup>) | 0.10 (0.43<sup>a</sup>) | 0.10 (0.43<sup>a</sup>) |
| $\langle I/\sigma I \rangle$ | 6.6 (2.0<sup>a</sup>) | 7.1 (2.0<sup>a</sup>) | 6.3 (2.0<sup>a</sup>) |
| average multiplicity | 2.0 (1.9<sup>a</sup>) | 2.3 (2.2<sup>a</sup>) | 2.0 (1.9<sup>a</sup>) |
| R-factor | 0.17 (0.28<sup>a</sup>) | 0.17 (0.31<sup>a</sup>) | 0.17 (0.27<sup>a</sup>) |
| R<sub>free</sub> | 0.22 (0.31<sup>a</sup>) | 0.23 (0.34<sup>a</sup>) | 0.22 (0.30<sup>a</sup>) |
| average B factor (Å<sup>2</sup>) | 33.7 | 34.9 | 30.2 |
| average B factor for inhibitor (Å<sup>2</sup>) | 65.9 | 72.4 | 58.8 |
| $\langle B \rangle$ from Wilson plot (Å<sup>2</sup>) | 59.9 | 19.5 | 40.6 |
| no. of protein atoms | 7710 | 7705 | 7695 |
| no. of inhibitor atoms | 96 | 80 | 84 |
| no. of solvent molecules | 699 | 488 | 645 |
| rms deviations from ideal values |  |  |  |
| bond lengths (Å) | 0.01 | 0.01 | 0.01 |
| bond angles (deg) | 1.65 | 1.57 | 1.61 |
| outliers in Ramachandran plot (%) | 0.30 | 0.60 | 0.40 |
| PDB code | 8Q18 | 8Q19 | 8Q1A |

<sup>a</sup>Values in parenthesis are for the high-resolution bin.

The electron density maps of the hCA IX-1, hCA IX-2, and hCA IX-3 complexes revealed interesting details about the binding modes of the saccharin-sulfonamide derivatives (Figure 3).

**Figure 3.**
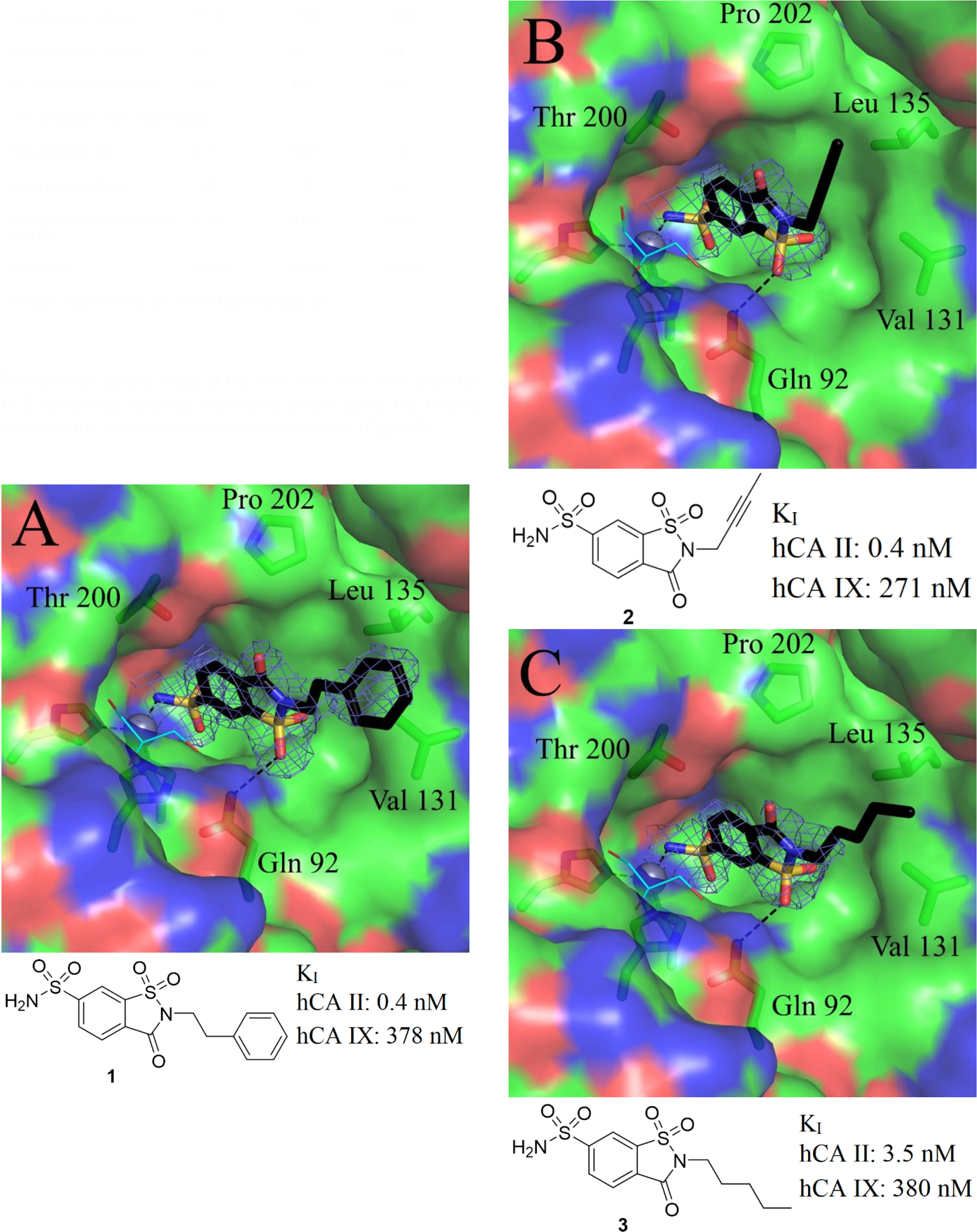
Comparison of the binding modes of compounds **1, 2**, and **3** within the active site of hCA IX. Panel A shows compound 1, panel B shows compound **2**, and panel C shows compound **3**. The protein is represented as a surface model, with surface atoms colored (carbon-green, nitrogen-blue, oxygen-red). The zinc ion is depicted as a gray sphere. Zinc coordinating histidines (His94, 96, and 119) and residues Gln92, Val131, Leu135, Thr200, and Pro202, which participate in hydrogen bonding, hydrophobic, and van der Waals contacts with the inhibitor, are indicated. A glycerol molecule bound to the enzyme is shown as a thin stick model with sky blue carbon atoms. For clarity, Fo-Fc OMIT electron density is shown only for the ligand and contoured at 3σ. The figure was prepared using PyMol (DeLano The PyMOL Molecular Graphics System San Carlos, CA, USA, DeLano Scientific).

In the hCA IX-1 complex, the electron density of the ligand was generally good, except for the benzyl moiety, which was weaker but still interpretable. Furthermore, a glycerol molecule was visible in the electron density map, bound within the active site of hCA IX in a nearly identical position to previously solved hCA IX structures.^[15]^ While the presence of the glycerol molecule during the cryo-protection of the crystal was assumed to have no effect on the binding and positioning of the inhibitor, its presence could potentially have an impact on the dynamics and energetics of the system. Similarly, in the hCA IX-2 and hCA IX-3 complexes, the electron density maps of the ligands were generally good, with the exception of the tail regions, where the electron density was much weaker. All three compounds exhibited similar binding patterns within the active site of hCA IX, with their tail moiety oriented towards a region that distinguishes it from other hCA isoforms. This region is considered a “hot spot” for designing isoform-selective hCA inhibitors. Surprisingly, despite differences in their chemical formulas and the absence of a hydrogen bond between the thiadiazole moiety of AAZ and Thr200, the binding orientations of these compounds in the hCA IX active site closely resembled that of the AAZ-hCA IX complex. The positioning of the ligands in the active site was facilitated by van der Waals and hydrophobic interactions involving Val131, Leu135, Thr200, and Pro202 residues (hCA II numbering), as depicted in Figure 4. Additionally, a hydrogen bond was observed between the ligands and Gln92, which is a common interaction seen in other sulfonamide-CAIs complexes.^[16,17]^

**Figure 4.**
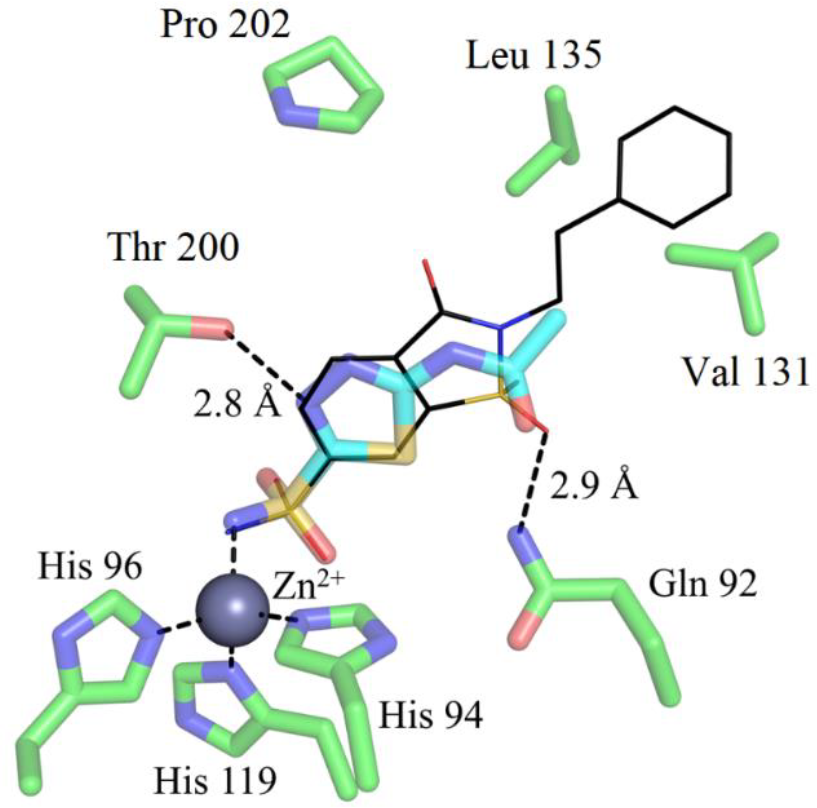
Comparison of binding modes of compound 1 (thin lines with black carbons), and CA inhibitor AAZ (PDB code 3IAI^[18]^, thick lines with green carbons) within the active site of hCA IX. The zinc ion is shown as a gray sphere and its coordinating histidines (His94, 96 and 119) are shown in green. Residues 92, 131, 135, 200 and 202 participating in hydrogen bonding, hydrophobic and van der Waals contacts with inhibitor are indicated. The figure prepared by using Pymol (DeLano The PyMOL Molecular Graphics System San Carlos, CA, USA, DeLano Scientific).

Our comparison of the hCA IX structures with previously published hCA II-saccharin derivative structures uncovered notable differences in their binding orientations (Figure 5).

**Figure 5.**
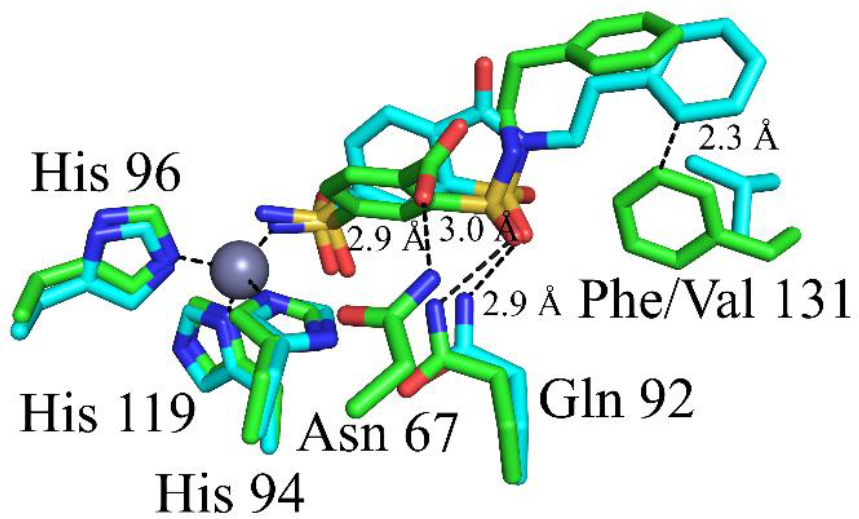
Comparison of binding modes of compound 1 within the active site of hCA IX (carbons are light blue) and hCA II (carbons are green, PDB code 5AMD^[10]^). The zinc ion is shown as a gray sphere and its coordinating histidines (His94, 96 and 119) and residues 67, 92 participating in hydrogen bonding with inhibitor are indicated. It should be noted that the orientation of the ligand’s tail in hCA IX differs from that in hCA II, possibly due to the potential clash with Phe131, as indicated in the figure. In hCA IX isoform ligands are bound without any changes in structure, while in hCA II active site, ligands were bound with open isothiazole ring, which probably occurred due alkaline hydrolysis. The figure was prepared using Pymol (DeLano The PyMOL Molecular Graphics System San Carlos, CA, USA, DeLano Scientific).

This observation highlights the importance of studying the interactions between potential drug candidates and different isoforms of hCA, as even small structural differences between isoforms can result in vastly different binding modes. Our findings provide valuable insights into the specific binding interactions and conformational changes that occur between hCA IX and the inhibitors. The fact that compounds **1, 2** and **3** exhibit different binding mechanisms between hCA II and hCA IX suggests the possibility of developing isoform-selective inhibitors for hCA IX, which could have significant implications in cancer treatment.

## Conclusion

The results of our study provide valuable insights into the binding modes of saccharin derivatives within the active site of hCA IX. Notably, compounds **1, 2** and **3** exhibit distinct binding modes when comparing hCA II and hCA IX. These compounds occupy a region within the active site that displays differences in both structure and sequence compared to other hCA isoforms. Through comprehensive characterization of their binding modes in the active sites of both hCA II and hCA IX, we have generated significant knowledge that can potentially be applied in the development of isoform-selective drugs targeting tumor-associated hCA IX.

## Experimental protocols

### Chemistry

The reports on the synthesis of the compounds can be found in the previous article.^[10]^

### CA inhibition assay

The inhibition data for saccharin derivatives 1-3 on hCA I, II, IX, and XII isoforms were obtained using the stopped-flow CO_2_ hydration method and are available in the previous article.^[10]^

### Protein production and purification

hCA IX catalytic domain (C174S, N346Q) was produced and purified in our lab as described earlier by our group. ^[15]^

### Crystallization

Protein was concentrated to 9 mg/mL in 20 mM Tris-HCl pH 8.0 using a 10 kDa cutoff Amicon concentrator. Crystallization was carried out by the sitting drop technique in 96-well MRC plates (Molecular Dimensions). For structure determination, 0.5 μl of protein was mixed with 0.5 μl of the bottom solution consisting of M di-ammonium hydrogen phosphate and 0.1 M sodium acetate pH 4.5. To fully grow crystals, 2 μl of mother liquor buffer containing 10 mM inhibitor was added and left for 5 days. The obtained crystals were soaked in mother liquor supplemented with 30% glycerol for approximately 1 min and then flash-frozen in liquid nitrogen.

### Data collection and structure determination

The datasets of hCA IX-ligand complexes were collected at BESSY II beamline 14.1 and processed using MOSFLM^[19]^ and SCALA^[20]^. Molecular replacement was performed using MOLREP^[21]^ with 5FL4^[15]^ as the initial model for hCA IX. Model refinement was conducted with REFMAC^[22]^ and the structures were visualized using COOT^[23]^. Ligand parameter files were generated using LIBCHECK^[24]^, and the ligand was manually fitted to the electron density map in COOT^[23]^. The coordinates and structure factors have been deposited in the PDB, and the PDB access codes, along with the data collection and refinement statistics, are shown in Table 2.

## Acknowledgements

This work was supported by the European Regional Development Fund (ERDF, for Jānis Leitāns project no. 1.1.1.2/VIAA/3/19/464; for Jekaterīna Ivanova project no. 1.1.1.2/VIAA/3/19/576).

## Entry for the Table of Contents

This study investigates the binding mechanisms of saccharine derivatives with human carbonic anhydrase IX (hCA IX), a vital antitumor target. Through crystallographic analysis, distinct binding modes are uncovered, revealing ligand-protein interactions pivotal for inhibitor efficacy and selectivity. This work enhances our understanding of hCA inhibitor binding and informs the rational design of potent agents.

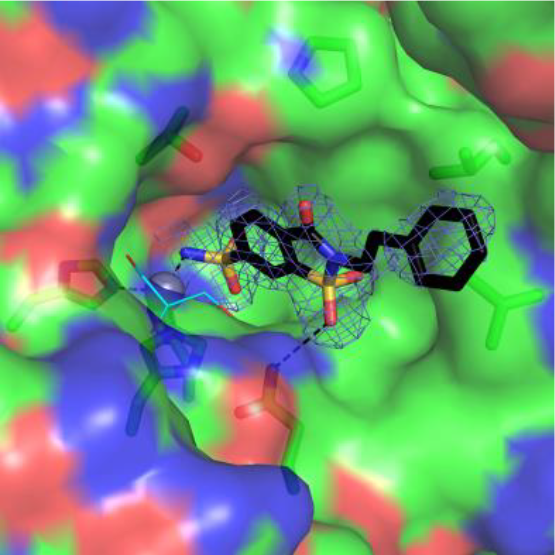

## Notes

### Competing Interest Statement

The authors have declared no competing interest.

